# Whole-genome sequences reveal zygotic composition in chimeric twins

**DOI:** 10.1101/2023.12.03.568663

**Authors:** Christopher J. Yoon, Chang Hyun Nam, Taewoo Kim, Jeong Seok Lee, Ryul Kim, Kijong Yi, June-Young Koh, Jiye Kim, Hyein Won, Ji Won Oh, Obi L. Griffith, Malachi Griffith, Joohon Sung, Tae Yeul Kim, Duck Cho, Ji Seon Choi, Young Seok Ju

## Abstract

The monochorionic placenta in dizygotic twins allows *in utero* exchange of embryonic cells, resulting in chimerism in the twins. In practice, this chimerism is incidentally identified on mixed ABO blood types or in the presence of cells with a discordant sex chromosome. Here, we applied whole-genome sequencing to one triplet and one twin families to precisely understand their zygotic compositions, using millions of genomic variants as barcodes of zygotic origins. Peripheral blood showed asymmetrical contributions from two sister zygotes, where one of the zygotes was the major clone in both twins. Single-cell RNA sequencing of peripheral blood tissues further showed differential contributions from the two sister zygotes across blood cell types. In contrast, buccal tissues were pure in genetic composition, suggesting that *in utero* cellular exchanges were confined to the blood tissues. Our study illustrates the cellular history of twinning during human development, which is critical for managing the health of chimeric individuals in the era of genomic medicine.

## Introduction

Sexually reproducing organisms begin as a single diploid zygote (2n) fertilized from a haploid sperm (n) and a haploid oocyte (n). The DNA from the zygote is then replicated as the embryo continues to divide and produce all the cells of an organism. In most individuals, every cell in an organism can be traced back to a single cell, the zygote. Chimeras are an exception to the norm in which an individual comprises cells with multiple genomic constitutions traced to more than one distinct diploid zygote (**Figure 1A**). In cattle, chimerism is frequently observed when multiple dizygotic bovine fetuses are fertilized^1,2^. Blood anastomosis in the monochorionic dizygotic (MCDZ) placenta allows the transfusion of cells and hormones between the two fetuses, often resulting in the generation of freemartin, an infertile female cattle. In humans, chimerism is rare as most human dizygotic twins have a dichorionic placenta that prohibits exchanges between the two twins^3^. Human chimeras are incidentally identified later in life when individuals show unusual phenotypes, such as sexual ambiguity^4^, lines of Blaschko^5^, abnormal blood typing results^6^, or incompatible parent-child relationships in genetic testing^7^.

**Figure 1.**
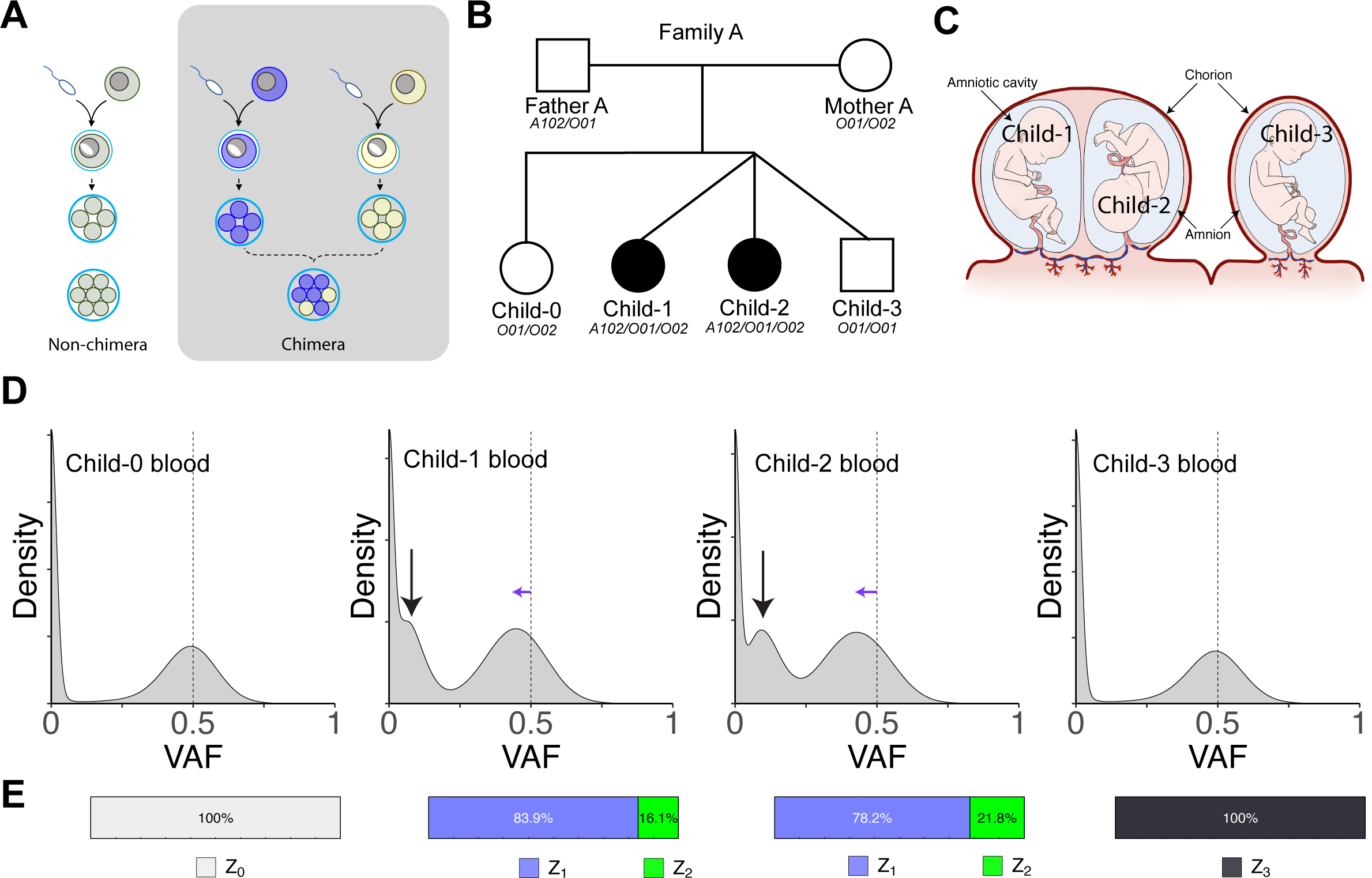
Quantification of hematopoietic chimerism in MCDZ twins. (A) Schematic of non-chimera and chimera. Non-chimeric individuals can be traced to one common zygote, while chimeric individuals are traced back to two zygotes. (B) A pedigree of Family A with chimera. Filled circles in Child-1 and Child-2 indicate the chimerism identified. ABO blood genotypes are shown below. (C) Fetal membranes (chorion and amnion) of the triplets in Family A. Child-1 and Child-2 showed monochorionic diamniotic configuration. Child-3 had a separate chorion and amnion. (D) VAF density of SNPs in the children. The dotted lines indicate VAF=0.5 expected for a heterozygous variant. Black vertical arrows indicate an extra peak shown in the chimeric individuals. The blue horizontal arrows indicate the left shift of the peak as a result of the merging heterozygous SNP peak (VAF=0.5) and an additional peak from chimeric SNPs. (E) The bar plots indicate the zygotic contribution estimated from the SNP read counts. MCDZ, monochorionic dizygotic; SNP, single nucleotide polymorphism; VAF, variant allele frequency

A few chimeric individuals have been reported in MCDZ twins^4,6,8–10^. Sex chromosomes^11–13^ or short tandem repeat markers^10–12^ have been utilized to evaluate the clonal composition of twins showing chimerism. However, these approaches lack the sensitivity and precision required to thoroughly assess the clonal composition of twins. The utilization of genome-wide variants can overcome these limitations and more comprehensively reveal the cellular history of chimerism. Thus, in the present study, we thoroughly investigated the inheritance patterns of millions of polymorphisms using whole-genome sequencing (WGS) and applied statistical evaluation to accurately decompose zygotic compositions at a single-cell and gamete resolution^14^. Furthermore, single-cell sequencing was used to trace the developmental outcomes of the chimeric cells in each twin.

## Results

### Chimerism in the blood of MCDZ twins

We explored the genome sequences of Family A (**Figure 1B**)^6^. Here, the first pregnancy gave birth to a healthy girl (Child-0). Then, the second pregnancy gave birth to triplets (Child-1, Child-2, and Child-3) who were conceived using assisted reproduction technology (ART) involving an ovulation induction with clomiphene citrate treatment. All triplet individuals were healthy, but two girls (Child-1 and Child-2) showed mixed ABO blood types (in the mixed-field agglutination of the blood groups). They had substantially dissimilar appearances, suggesting a non-identical genetic constitution. The triplets were noted to have a dichorionic triamniotic placenta (**Figure 1C**) where Child-1 and Child-2 shared a single chorion (i.e., a monochorionic diamniotic configuration between Child-1 and Child-2).

To systematically understand chimerism, we quantified the level of chimerism in the peripheral blood using the WGS of the family members. Here, we investigated the variant allele frequency (VAF) of inherited single-nucleotide polymorphisms (SNPs), in which one of the parents had a heterozygous genotype, and the other parent had a homozygous reference genotype. Under Mendelian inheritance^14^, an offspring genome will have SNPs that are either heterozygous genotypes (VAF∼0.5) or homozygous reference genotypes (VAF∼0) with a 50%:50% chance. Suppose the genomes of two different zygotes are admixed; intermediate VAFs (between 0<VAF<0.5 according to the mixture level) will be observed in the genomic regions where the two zygotes inherited different parental chromatids.

The peripheral blood of Child-0 and Child-3 showed a typical VAF pattern of the non-chimeric genome, or single zygote, with two possible genotypes (VAF∼0; homozygous reference genotype, VAF∼0.5; heterozygous genotype; **Figure 1D**). In contrast, the peripheral blood of Child-1 and Child-2 showed additional peaks near VAF∼0.1 in Child-1 and Child-2 and left shifted peak near VAF∼0.4, indicating chimerism in their blood, implying the presence of admixed cells of dual zygotic origins in the blood of both twins (**Figure 1D**).

One of the possible sources of the chimerism is the male sibling (Child-3) in the triplet pregnancy. However, this hypothesis was rejected since Child-3 specific variants and chromosome Y were not observed in the blood DNA of Child-1 and Child-2 (**Supplemental Figure S1A**). Instead, a comparison of the SNPs in the blood tissues of Child-1 and Child-2 suggested that the chimerisms of these twins were associated with a mutual exchange of cells from two zygotic lineages, independent to Child-3.

Indeed, the genomic sequences of peripheral blood tissues of Child-1 and Child-2 were successfully decomposed by two sets of zygotes (Z_1_ and Z_2_; **Figure 1E**). A maximum likelihood estimation approach was used to quantify the genomic compositions of the two sibling zygotes^14^ (Z_1_:Z_2_ = 78.2%:21.8% and 83.9%:16.1% for Child-1 and Child-2, respectively, **Figure 1E**). We investigated another MCDZ twin family (Family B, **Supplemental Figures S2A, S2B**) showing a similar pattern of zygotic contributions (Z_4_: Z_5_ = 91.2%:8.8% in Child-21 and 90.0%: 10.0% in Child-22, **Supplemental Figure S2C**). In both the MCDZ twins, one of the sister zygotes was commonly dominant in the blood. This indicated that immigrant cells became a predominant population in the blood tissues of both the twins.

### Decomposition of haplotype-resolved zygote-specific genome sequence in the chimera

Genomic sequences of the parents and multiple zygotes (Z_0_, Z_1_, Z_2_, and Z_3_, where Z_0_ and Z_3_ are zygotes generating Child-0 and Child-3, respectively) in the family allowed us to phase the haplotype genome sequences in the parents and their meiotic recombination breakpoints in each zygote^15^ (**Figure 2A**). The genomic configurations of Child-1 and Child-2 can be categorized into three groups (**Supplemental Figure S3A**): regions with two haplotypes (Z_1_ and Z_2_ receiving identical chromosome segments from both parents), three haplotypes (both zygotes receiving identical chromosome segments from one of the parents, but receiving opposite homologous chromosomes from the other parent), or four haplotypes (both zygotes receiving opposite chromosome segments from both parents), according to the meiotic recombination and chromosomal segregation stochastically operative in the gametogenesis. Overall, the entire autosomal genome of Child-1 and Child-2 contained 26.6% two-chromosome regions, 46.2% three-chromosome regions, and 27.2% four-chromosome regions (**Supplemental Figure S3B**), which was close to the random expectation (25%:50%:25%; chi-squared test P-value=0.9971). Since the two-haplotype regions represent the identical sequence between the two zygotes, the remaining 73.4% (three and four-haplotype regions) of the autosomal genome were in a true chimeric state.

**Figure 2.**
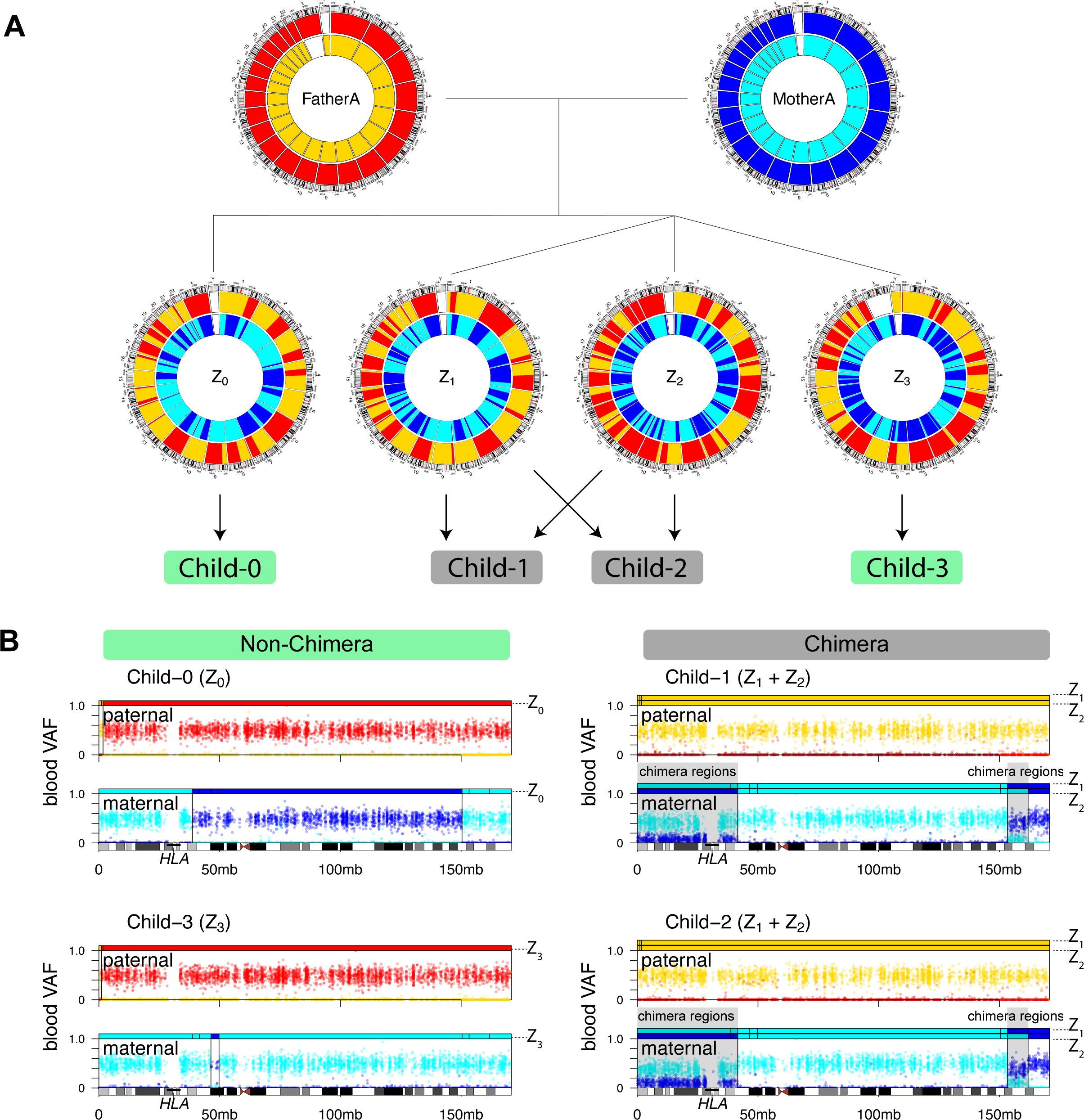
Haplotype resolved diploid genomes of blood chimerism. (A) A circos diagram of the parents and the four zygotes in Family A. Each haploid of parental chromosomes is assigned specific colors, and meiotic recombinations are seen as a color switch within the chromosomes of the zygotes. Contributing zygote(s) for each child are indicated with arrows. (B) VAFs of each child are shown with their respective parental haploid of origin as their color (same color as Figure 2A). The parental haploid genomes are also shown as rectangles over each plot. Two bars are drawn for chimeric children to indicate two originating zygotes. The regions where the zygotes inherited the same haplotype block are shown identical colors with VAF∼0.5. The regions where the zygotes inherited different haplotype blocks are highlighted with a gray background. VAF, variant allele frequency

Several genes of interest were located in the chimeric regions. For example, the *HLA* genes were located in a three-chromosome region (chromosome 6), where two sets of maternal chromosomes contributed to chimerism (**Figure 2B**). The *ABO* gene was located in a three-chromosome region (chromosome 9), where zygote Z_1_ and Z_2_ inherited different chromosomal segments from the father (Z_1_: *O01* allele; Z_2_: *A102* allele), but the same chromosome segment from the mother (*O02* allele for both Z_1_ and Z_2_), fully explaining the ambiguous blood type observed in Child-1 and Child-2 (**Supplemental Figure S1B**).

### Contribution of each zygote in various blood cell types

Inherited SNPs unique to Z_1_ and Z_2_ can be used as molecular markers to differentiate the zygotic origins of each single cell in Child-1 and Child-2. We decomposed the zygotic origin of peripheral blood mononuclear cells (PBMCs) in the chimeric twins. To this end, PBMCs of Child-1 and Child-2 were profiled by single-cell transcriptome sequencing, then were clustered into various cell types, including CD4^+^ T lymphocytes, CD8^+^ T lymphocytes, B lymphocytes, natural killer (NK) cells, and monocytes (**Figure 3A)**. The zygotic origins of these cells were further assigned using their genotype information from the single-cell sequencing data (**Figure 3B**)^16^. Although the dominance of the zygote (Z_1_) was consistent in all cell types, the relative fraction between Z_1_ and Z_2_ differed for each cell type (**Figure 3C**; P < 0.05, Two-proportion Z-test). Notably, the proportions of Z_2_ were higher in CD4^+^ T cells and NK cells but lower in monocytes in both the twin individuals, suggesting a genetic fitness of the two zygotes into the relative differentiation potential to a specific cell type.

**Figure 3.**
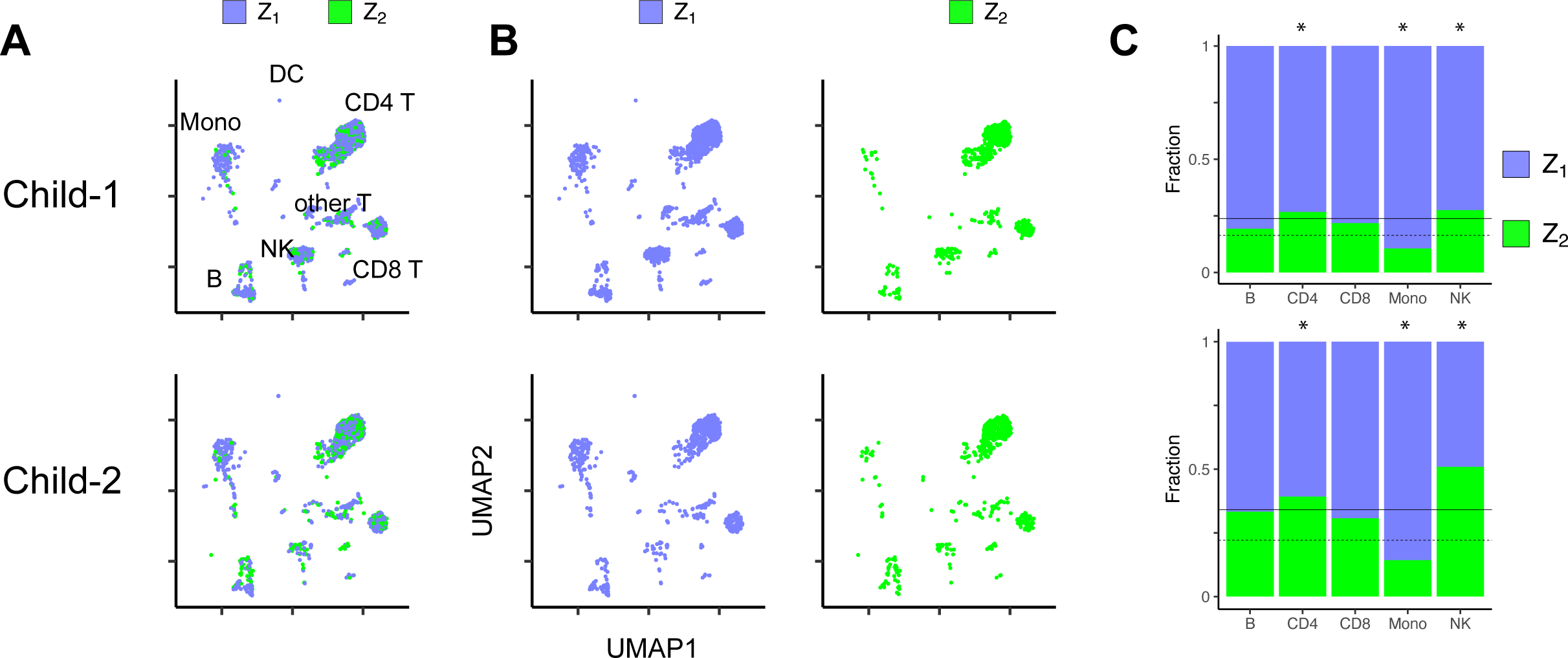
Single-cell resolved chimerism of peripheral blood mononuclear cells. (A) UMAP projection of the PBMC of Child-1 and Child-2 in Family A. (B) Each contributing zygote is drawn separately for Child-1 and Child-2. (C) Cell type-specific zygotic fractions in the PBMCs of Child-1 and Child-2. The asterisks indicate statistically significant differences compared to overall cell counts (two-proportion Z-test, *P<0.05). PBMC, peripheral blood mononuclear cell; UMAP, uniform manifold approximation and projection

### Tracing the developmental history of chimerism in non-hematopoietic tissues

To explore potential chimerism in non-blood solid tissues, we sequenced the whole-genome of the buccal tissues of the twin individuals (Child-1 and Child-2). As buccal tissues are usually contaminated by blood cells to a substantial level^17^, we carefully isolated the buccal epithelial cells using the laser capture microdissection technique (LCM)^18^(**Figure 4A**). In contrast to the blood tissues, we found no supporting evidence of chimerism in LCM-isolated pure buccal epithelial cells (**Figure 4B**). Among LCM-isolated buccal epithelial cells from Child-1 (n=325 cells), we did not observe any Z_2_ originating epithelial cells (**Supplemental Figure S4**). Conversely, among LCM-isolated buccal epithelial cells from Child2 (n=67 cells), no Z_1_ originating epithelial cells were observed (**Supplemental Figure S4)**. Our data indicated that cellular exchanges are likely confined to hematopoietic stem cells. Our results indicate that the dominant zygotes are discordant across the tissues in Child-2, i.e., Z_1_ for blood cells and Z_2_ for buccal epithelial cells and potentially other solid tissues as well.

**Figure 4.**
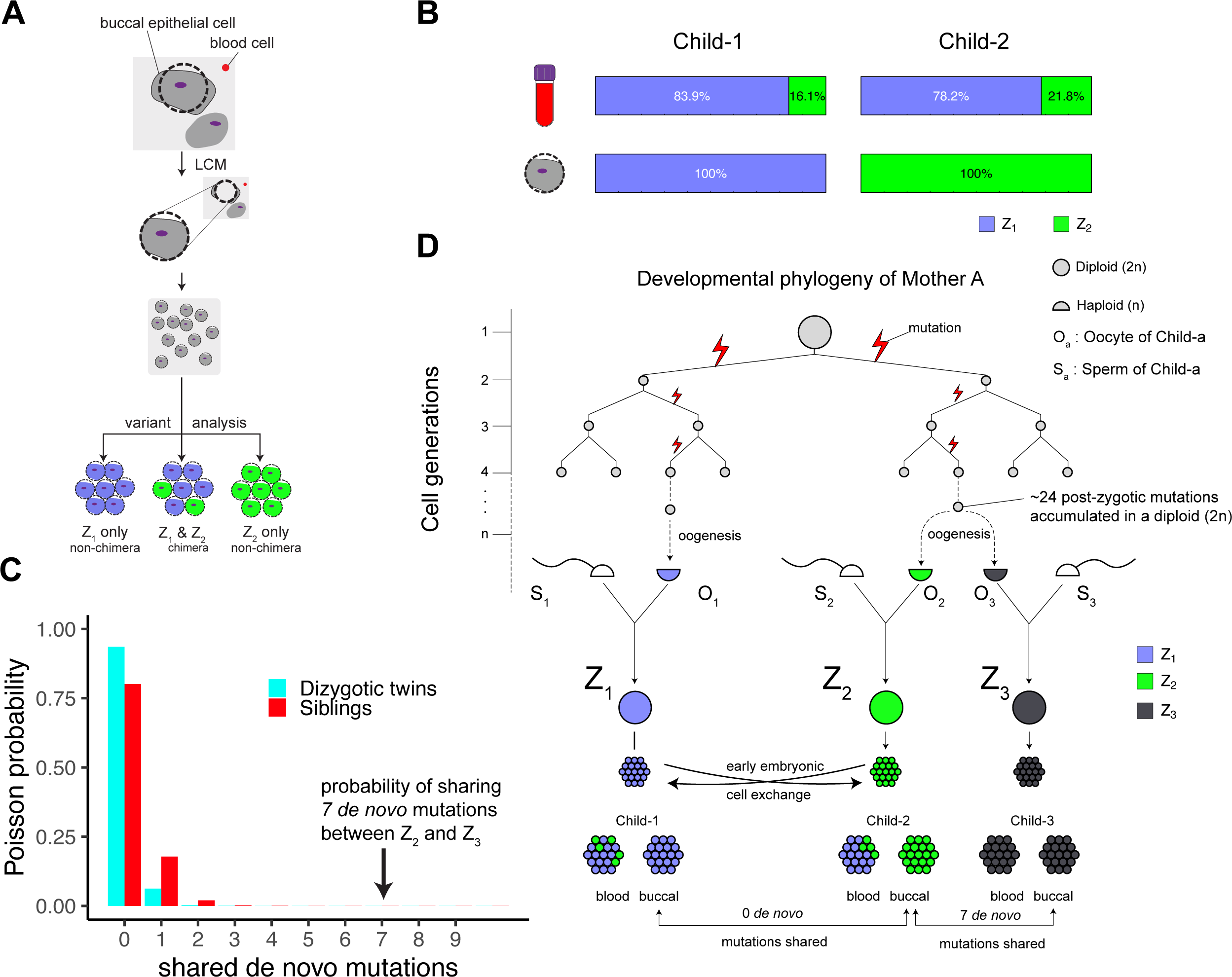
Tracing the chimerism of MCDZ chimera in non-hematopoietic tissues. (A) Single buccal epithelial cells are isolated using laser capture microdissection (LCM). Isolated buccal epithelial cells are pooled to generate sequence libraries. Variant analysis of each sequencing library was used to determine the contribution of the zygotes in the buccal epithelial cells. (B) The contributions of f Z1 and Z2 were estimated in Child-1 and Child-2’s peripheral blood (top) and LCM-isolated buccal epithelial cells (bottom). The zygotic contributions for peripheral blood (top) for Child-1 and Child-2 are identical to Figure 1E, shown for comparison with those in buccal epithelial cells. (C) The height of each bar represents the Poisson probability of shared *de novo* mutation counts based on reported data from dizygotic twins and siblings^21^. The number of shared *de novo* mutations (7) between Z2 and Z3 is shown with an arrow. (D) The lineage history of the triplets is traced to gametogenesis. The half circles indicate haploid genomes of sperm or oocytes. The full circles indicate diploid genomes in the fertilized eggs and embryonic cells. MCDZ, monochorionic dizygotic

### Potential impact of ovulation induction in oogenesis

Every individual typically has approximately 70 (50–100) *de novo* mutations, which generate during parental gametogenesis^19,20^. On average, approximately 20% of *de novo* mutations are of maternal origin^19,20^. Usually, a sibling pair rarely shares *de novo* mutations because the chance of selecting two gametes sharing a close cellular lineage (which will share *de novo* mutations) within a large cellular pool of gametes is extremely low^21^. Indeed, sequencing data from the two external cohorts showed that most sibling pairs shared 0 or 1 *de novo* mutation (**Figure 4C**).

Unexpectedly, we observed an unusually high number of shared *de novo* mutations between Z_2_ (one of the zygotes contributing to Child-1 and Child-2) and Z_3_ (the zygote contributing to Child-3) (n=7; P = 4.37×10^-9^, exact Poisson test λ=0.22, 95% CI, 2.8 to 14.42, **Figure 4C, Supplemental Table S1**). We speculated that all these shared *de novo* mutations were of maternal origins, as they were all located in the genomic regions of the identical maternal haplotypes in Z_2_ and Z_3_ (**Figure 2A**). Further, the two informative shared *de novo* mutations (chr3:195,327,945 A>T and chrX:10,147,343 C>T) were directly phased to the nearby maternal germline polymorphisms, confirming their maternal inheritance. In a thought experiment in which the same primary oocyte iteratively generated mature egg cells, a random pair of these egg cells shared seven *de novo* mutations (70 *de novo* mutations × 20% of maternal origins × 50% shared haplotypes), which is very close to the observed number of shared *de novo* mutations. Therefore, our observations imply that the oocytes contributing to Z_2_ and Z_3_ diverged very recently from the maternal germline (**Figure 4D**). Given the shared maternal haplotypes between Z_2_ and Z_3_ (56.9%), we speculated that the most recent common ancestor cell of maternal lines of the zygotes harbored 24 post-zygotic mutations (approximately a quarter of which are inherited as *de novo* mutation in the offspring) in the diploid genome. The mutation burden was even higher to the expected number of mutations in a diploid primary oocyte (n = 10-20), implying that two oocytes for Z_2_ and Z_3_, were branched very recently in Mother A (**Figure 4D**). Even though it is an observation from one family, we cannot rule out the possibility that ovulation induction treatment may have stimulated oogenesis and ovulation from a particular gamete lineage, contributing to two of the three fertilized oocytes in the triplet pregnancy in Family A.

## Discussion

In this study, we used genome sequencing technologies to understand zygotic composition and to further infer the developmental history of chimerism in human MCDZ twins. We used a WGS-based approach using millions of SNPs scattered throughout the genome, to quantify the contribution of each zygote in the chimeric twins. In both the MCDZ twin families investigated in this study, there was a clear dominant zygote in the blood of the twins, likely resulting from an early extensive exchange and engraftment of hematopoietic stem cells. A long-term follow-up of these individuals is be necessary to understand whether their chimeric ratios are stable over time throughout their lives.

Although it is unclear whether chimerism occurs beyond the blood tissues of MCDZ twins, this study indicates that chimerism can be restricted to the hematopoietic lineages. However, our investigation should be interpreted carefully as it was limited to only two tissues (buccal epithelial and blood cells) from a twin pair. A study on additional cell types is needed to determine whether the chimerism is truly restricted to the hematopoietic system or is present in other lineages. A long-term follow-up study will be helpful in collecting other cell types from these individuals in case of medical procedures, such as surgery and/or biopsy.

The chimeric children are currently healthy, but the clinical significance of chimerism remains unknown. Our observations directly indicate that two sets of antigen-presenting cells, having haploidentical HLA types, and two sets of lymphocytes are present in each twin individual. Long-term investigation of the immunological phenotypes of these twin individuals will be also helpful for understanding the regulation of immune tolerance in chimeric individuals.

Finally, our analysis revealed an unexpectedly close relationship at the gamete level in the twins. We speculate that ovulation induction is responsible for the close genetic lineages, and further large-scale analyses of twins or multiple births from ovulation induction are required to prove the hypothesis.

In this work, we present the most detailed life history of human chimerism in MCDZ twins to date. The study on MCDZ cattle (freemartin) led to the Nobel Prize winning discovery of acquired immune tolerance, suggesting the idea that immunity is established during embryogenesis^22^. Little is known about the life history of chimerism and immunological consequences of human MCDZ twins. As ART is being utilized more frequently to aid in human reproduction, the possibility of the formation of MCDZ twins and shared *de novo* mutations among siblings must be assessed to provide appropriate medical care for these individuals in the era of personalized and genomic medicine.

## Supporting information

Key resources table

## Acknowledgments

This work was supported by the National Institutes of Health MSTP grant (T32 GM07200 to C.J.Y.) and the National Institutes of Health NRSA Predoctoral Fellowship (F30 HD106744 to C.J.Y.). This work was supported by the National Research Foundation of Korea (NRF) grant funded by the Korea government, Ministry of Science and ICT (No. NRF-2022R1A5A102641311 to Y.S.J; Leading Researcher Program NRF-2020R1A3B2078973 to Y.S.J; to NRF-2020M3A9E4039653 to Y.S.J).

## Author contributions

Conceptualization, C.J.Y. and Y.S.J.; Methodology, C.J.Y.; Investigation, C.J.Y., J.S.L., K.Y., J-Y.K., R.K., C.H.N., J.K., T.K., H.W., J.W.O., Y.S.J.; Formal analysis, C.J.Y.; Writing – Original Draft, C.J.Y.; Writing – Review & Editing, C.J.Y., Y.S.J. Funding Acquisition, C.J.Y. and Y.S.J.; Resources, J.S.C., T.Y.K, D.C., J.S.; Supervision, O.L.G., M.G., and Y.S.J.

## STAR METHODS

## RESOURCE AVAILABILITY

### Lead Contact

Further information and requests for resources and reagents should be directed to and will be fulfilled by the Lead Contact, Young Seok Ju

### Material availability

This study did not generate new unique reagents.

### Data and code availability

The blood and buccal WGS data of both families can be accessed at the European Nucleotide Archive under the accession number EGAS00001005997. LCM-isolated buccal cell low-input WGS data and single-cell RNA sequencing data are available under the accession number EGAS00001007649. All original code has been deposited at Zenodo (https://doi.org/10.5281/zenodo.10151688) and is publicly available as of the publication date. The DOI is listed in the key resources table. Any additional information required to reanalyze the data reported in this paper is available from the lead contact upon request.

## STUDY PARTICIPANT DETAILS

### Study participants

Two families with known chimerism identified by ABO genotyping were recruited for the study^6^. In the first pregnancy, a health girl (Child-0) was conceived. The second pregnancy gave birth to triplets (Child-1, Child-2, and Child-3). The second pregnancy was conceived using ART, involving ovulation induction with clomiphene citrate treatment. Peripheral blood samples were obtained for all individuals. Buccal swab samples were also obtained from chimeric twins in each family. Child-1 and Child-2 were five years old at the time of whole-genome sequencing. Child-1 and Child-2 were six years old at the time of peripheral blood mononuclear cell (PBMC) and buccal epithelial LCM isolation studies.

### Ethics approval and consent to participate

The study protocol was approved by the Institutional Review Boards of the Korea Advanced Institute of Science and Technology (IRB KH2019-174), International St. Mary’s Hospital (IRB IS19TIME0070), and the Samsung Medical Center (IRB SMC-2019-01-049-005).

## METHOD DETAILS

### Whole-genome sequencing (WGS) data generation

DNA was extracted from the peripheral blood samples of all family members. Buccal swabs were obtained from the chimeric individuals. DNA libraries were generated according to the Illumina’s Truseq PCR free library protocol. Paired-end sequencing reads of WGS were obtained using NovaSeq (blood: 30x, buccal: 60x). Raw sequence files were aligned to the human reference genome GRCh38 using the BWA mem^23^.

### Investigation of inherited heterozygous loci

Germline single nucleotide polymorphisms (SNPs) and small indels were used to estimate the level of chimerism in the children. Briefly, for an informative locus where one of the parents was a homozygous reference (ref/ref) and the other was a heterozygous variant (ref/var), there were two possible genotypes in the daughter embryo (homozygous reference and heterozygous variant). In genetically homogeneous offspring, the variant allele fraction (VAFs) of the locus in sequencing is either 0% (ref/ref) or 50% (ref/var). In contrast, for chimeric offspring homozygous reference and heterozygous variant cells co-existed in a single individual, leading to a dispersion of VAF between 0 and 50%. We used TrioMix, a maximum likelihood estimation framework, to quantify chimerism in the twins using the inheritance pattern of common SNPs in GRCh38^14^.

### Meiotic recombination

Meiotic recombination was identified by tracing the parental haplotype in the non-chimeric siblings (Child-0 and Child-3) and the monochorionic twin sisters (Child-1 and Child-2). The circular Binary Segmentation method^26^ was used to estimate the recombination sites. All recombination sites were manually reviewed, and parental haplotypes were inferred from the recombination patterns using the most parsimonious recombination breakpoints between the siblings^15^.

### Non-inherited variant discovery

Non-inherited variants (*de novo* mutations and postzygotic mutations) were identified using Varscan2^24^ and in-house scripts from our previous reports^25^. Non-inherited variants were compared between Child-0, Child-1, Child-2, and Child-3. All *de novo* mutations shared between the children were manually reviewed and compared against meiosis recombination patterns to infer the parental origin of shared *de novo* mutations.

### Laser-capture microdissection (LCM) and low input library preparation

Cells obtained from the buccal swab were placed on membrane-covered microscope slides (Carl Zeiss Membrane Slides). Cells on the microscope slides were fixed using a PAXgene fixation kit, followed by hematoxylin and eosin staining and 70% alcohol treatment. Slides were mounted tono an LCM machine (Carl Zeiss, PALM MicroBeam). Single buccal epithelial cells were dissected and collected on an adhesive cap (Carl Zeiss Adhesive Caps) with approximately 13–60 cells per cap. Each adhesive cap at an density of approximately 13–60 cells was independently processed for DNA extraction using a QIAamp DNA micro kit (Qiagen). This approach allowed enough DNA input material to prepare sequencing libraries and detect variants unique to each zygote. THe NEBNext Ultra II FS DNA library kit was used to prepare low-input DNA libraries. Paired-end whole genome sequencing reads were obtained using a NovaSeq machine as described above.

### 10X single-cell transcriptome sequencing

PBMCs were isolated from the whole blood using Ficoll density gradient centrifugation. PBMC of the chimeric children were then prepared using a 10x genomics 3’ gene expression kit for single cell RNA sequencing. 10x single cell libraries were sequenced using the Illumina NovaSeq. Demultiplexed FASTQ files were processed using CellRanger (v6.0.1). Output matrix was used as the input for Seurat (v4) for clustering and cell type annotation. Cell types were mapped to a two dimensional uniform manifold approximation and projection space of multimodal reference atlas of circulating human immune cells^27^.

### Phasing variants to each zygote and deconvolution of single-cell sequencing data

The zygote of origin of each cell was determined using a custom script utilizing high-confidence SNP locations specific to one of the two contributing zygotes in the chimera. High-confidence SNP loci were obtained by comparing the SNP VAFs in the buccal swab whole-genome sequences with each zygote’s recombination patterns. Variants were phased to one of the parental homologous chromosomes using variant information from the additional siblings (Child-0 and Child-3). We could accurately assign their genotypes because the additional siblings were non-chimeric individuals. Observation of the variant read counts follows a binomial probability (Pr) distribution where *k* is an alternative read count, *n* is a total read depth at a locus, and *α* is the probability of sampling an alternative read, which is identical to the zygote fraction for homozygous variants.

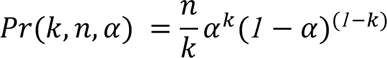

Cumulative distribution function *F*(*k*, *n*, *α*) provides a confidence interval for observing the read counts for a given *α*.

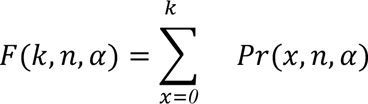

For a heterozygous variant, *α* is 0.5. Thus, a variant wasconsidered a heterozygous variant with a 95% confidence under binomial probability if it satisfied the following condition:

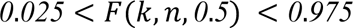

We required VAF=0 for a homozygous reference genotype and the observed variant read counts to satisfy the above equation. For Child-0 and Child-3 we compared the genotypes of the two siblings and the chromosomal phasing information to assign each variant to its respective homologous chromosomes in the parents.

For the two zygotes to have a homozygous variant that differs in genotype (one zygote with a homozygous alternative genotype and the other zygote with a homozygous reference genotype), the two zygotes must receive the opposite allele from both parents which is only possible in the four-chromosome region. Let us assume that Z_1_ and Z_2_ fraction is *α_1_* and 1-*α_1_* respectively in Child-1’s bulk buccal WGS and 1-*α_2_*and *α_2_*in Child-2’s bulk buccal WGS. A variant locus *j* from the bulk buccal WGS of Child-1 was observed with a total read depth *n*_’_*_1_*and an alternative read count *k*_’_*_1_*and Child-2 with a total read depth *n*_’_*_2_* and a read count *k*_’_*_2_*. To identify the homozygous variant *j* specific to Z_1_, but not Z_2_, with a 95% confidence interval, the read counts from the two bulk buccal WGS must satisfy the following conditions.

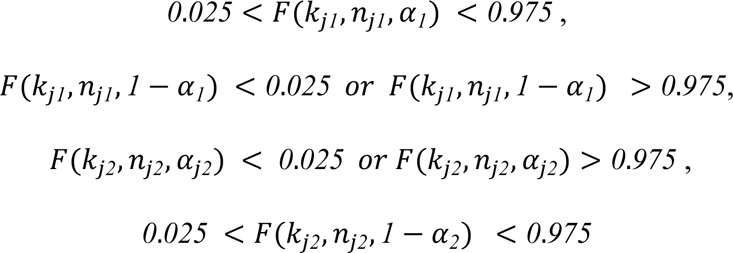

Using these statistical assignments of unique variants, we obtained 44,448 homozygous variants unique to Z_1_ and 44,193 homozygous variants unique to Z_2_. Homozygous variants specific to Z_2_, but not Z_1,_ could be similarly calculated.

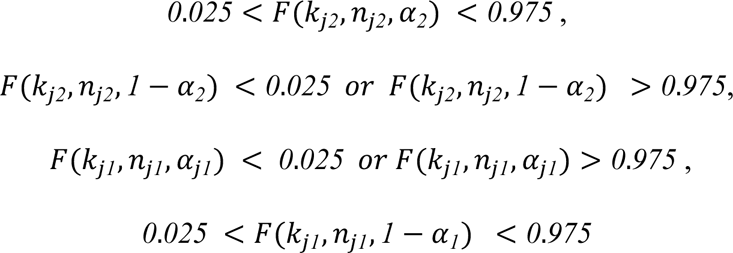

For heterozygous variants specific to each zygote, we substitute *α* and 1-*α* with 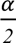 and 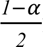 for the alternative allele frequency. We obtained 462,339 heterozygous variants specific to Z_1_ and 456,409 heterozygous variants specific to Z_2_.

A VCF file containing the zygote-specific heterozygous and homozygous SNPs was used as an input for Demuxlet^16^ to identify singlet cell barcodes with their zygotes of origin identity. Cell barcodes containing SNPs from both zygotes were labeled doublets or multiplets and removed from further analysis.

## QUANTIFICATION AND STATISTICAL ANALYSIS

Maximum likelihood estimation was used to estimate the degree of chimerism using TrioMix by quantifying the deviation from the expected Mendelian inheritance patterns of the SNPs^14^. Linear regression was performed between the expected ratio of Z_1_ and Z_2_ and the bulk sequencing VAFs of the mitochondrial heteroplasmic variants. Chi-squared test was used to estimate the two-, three, and four-chromosome regions of the chimera. A two-proportion Z-test was used to estimate the differential distribution of zygotic origins in various cell types using scRNA-seq. Poisson’s probabilities were used to estimate the likelihood of observing shared *de novo* mutations in genetic sibling zygotes. All statistical calculations and visualizations were conducted using R (v4.0) programming language.

## SUPPLEMENTAL INFORMATION

**Table S1. De novo mutation shared between the Z2 and Z3.**

**Figure S1. Whole-genome sequence reads of peripheral blood of Family A.**

**Figure S2. Quantification of the chimerism in Family B**

**Figure S3. Chromosome distribution of the two zygotes composing the chimera twins**

**Figure S4. Single-cell isolated buccal epithelial cell mitochondrial DNA variant analysis**

---

**Table S1.** De novo mutation shared between Z_2_ and Z_3_. The buccal epithelial cells of Child-2 were used as a proxy for Z_2_ (Child-2 buccal WGS estimated to be 98.11% of Z_2_). Raw read counts and VAFs were calculated for the Child-2 and Child-3 buccal WGS. The phasing column indicates read-level phasing results of the variant to maternally inherited SNPs. SNP, single nucleotide polymorphism; WGS, whole-genome sequencing; VAF, variant allele frequency

| Position | Rer base | Alt base | Buccal<br>Child2 ALT | Buccal<br>Child2 Depth | Buccal<br>Child2 VAF | Buccal<br>Child3 ALT | Buccal<br>Child3 Depth | Buccal<br>Child3 VAF | Phasing |
| --- | --- | --- | --- | --- | --- | --- | --- | --- | --- |
| chr10:56883013 | T | A | 19 | 64 | 0.297 | 18 | 33 | 0.546 | FALSE |
| chr12:86289509 | TAAC | T | 19 | 54 | 0.352 | 17 | 34 | 0.500 | FALSE |
| chr22:47996473 | C | T | 30 | 70 | 0.429 | 22 | 42 | 0.524 | FALSE |
| chr2:30355496 | G | T | 29 | 70 | 0.414 | 13 | 24 | 0.542 | FALSE |
| chr3:184936466 | C | T | 25 | 47 | 0.532 | 12 | 28 | 0.429 | FALSE |
| chr3:195327945 | A | T | 24 | 54 | 0.444 | 8 | 18 | 0.444 | TRUE |
| chrX:10147343 | C | T | 33 | 65 | 0.508 | 11 | 11 | 1.000 | TRUE |

