## Supplemental figures for "Whole-genome sequences reveal zygotic composition in chimeric twins"

### Supplementary Data

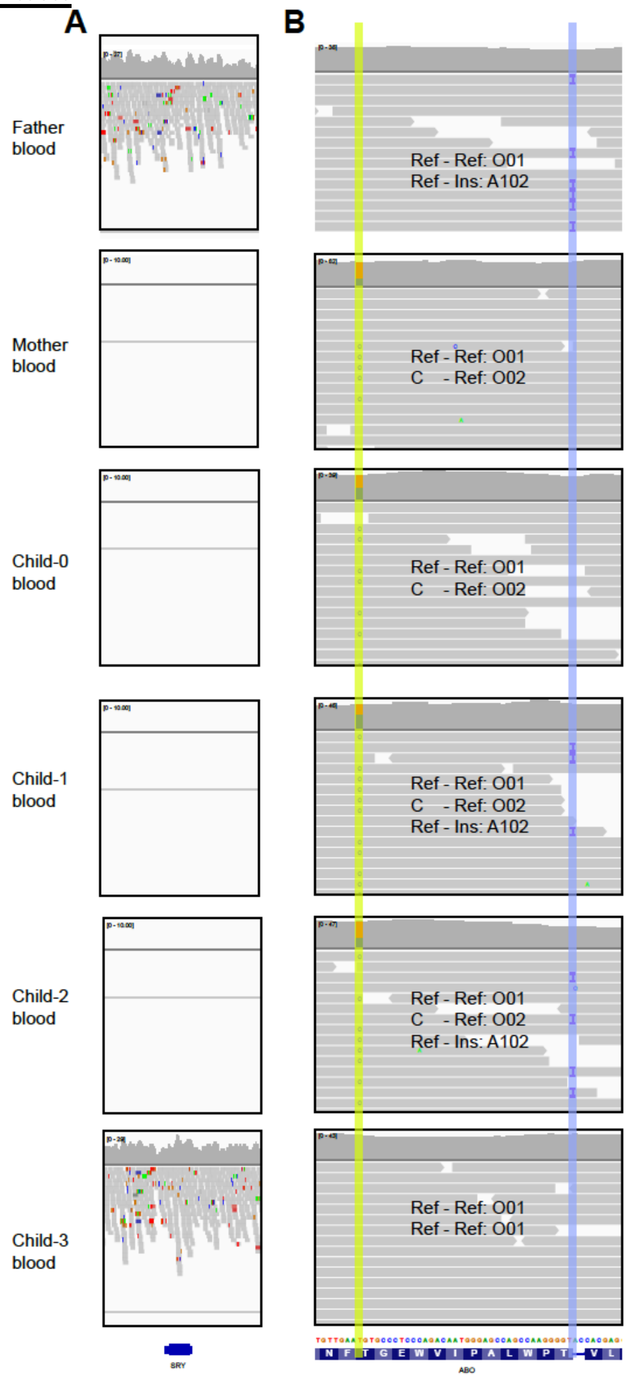


**Supplemental Figure S1**. **Whole-genome sequence reads of peripheral blood of Family A. (A)** The region shown is the *SRY* gene locus in chromosome Y. (B) Read-level genotyping of the *ABO* gene locus is shown in the IGV screenshot. Three haplotype groups are seen in Child-1 and Child-2, unlike non-chimeric individuals. SNV determining the O02 genotype is highlighted in yellow, while the indel determining the O genotype vs A102 is highlighted in blue. All samples shown are from blood DNA.

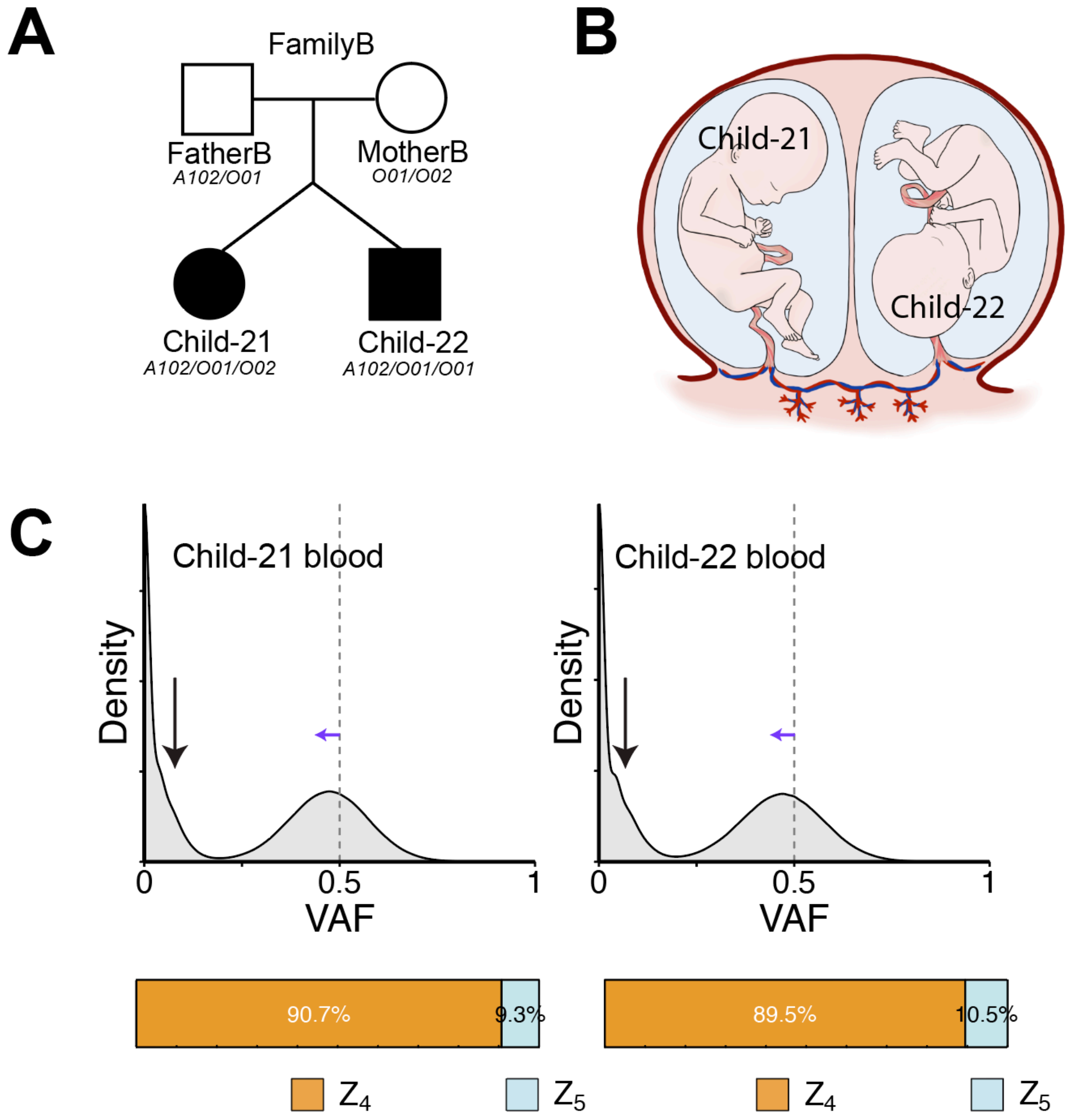


**Supplemental Fig S2. Quantification of chimerism in Family B.** (A) Pedigree of FamilyB. The two children in Family B were chimera and indicated with filled circles. Blood genotypes of all individuals are shown below. (B) Fetal membranes (chorion and amnion) of the twins in FamilyB. Child-21 and Child-22 showed monochorionic diamniotic configuration. (C) VAF density plot for blood DNA of Child-21 and Child-22. The dotted lines indicate VAF=0.5 expected for a heterozygous variant. The black vertical arrows indicate an extra peak shown in the chimeric individuals. The blue horizontal arrows indicate the left shift of the peak as a result of the merging heterozygous SNP peak (VAF=0.5) and an additional peak from chimeric SNPs.

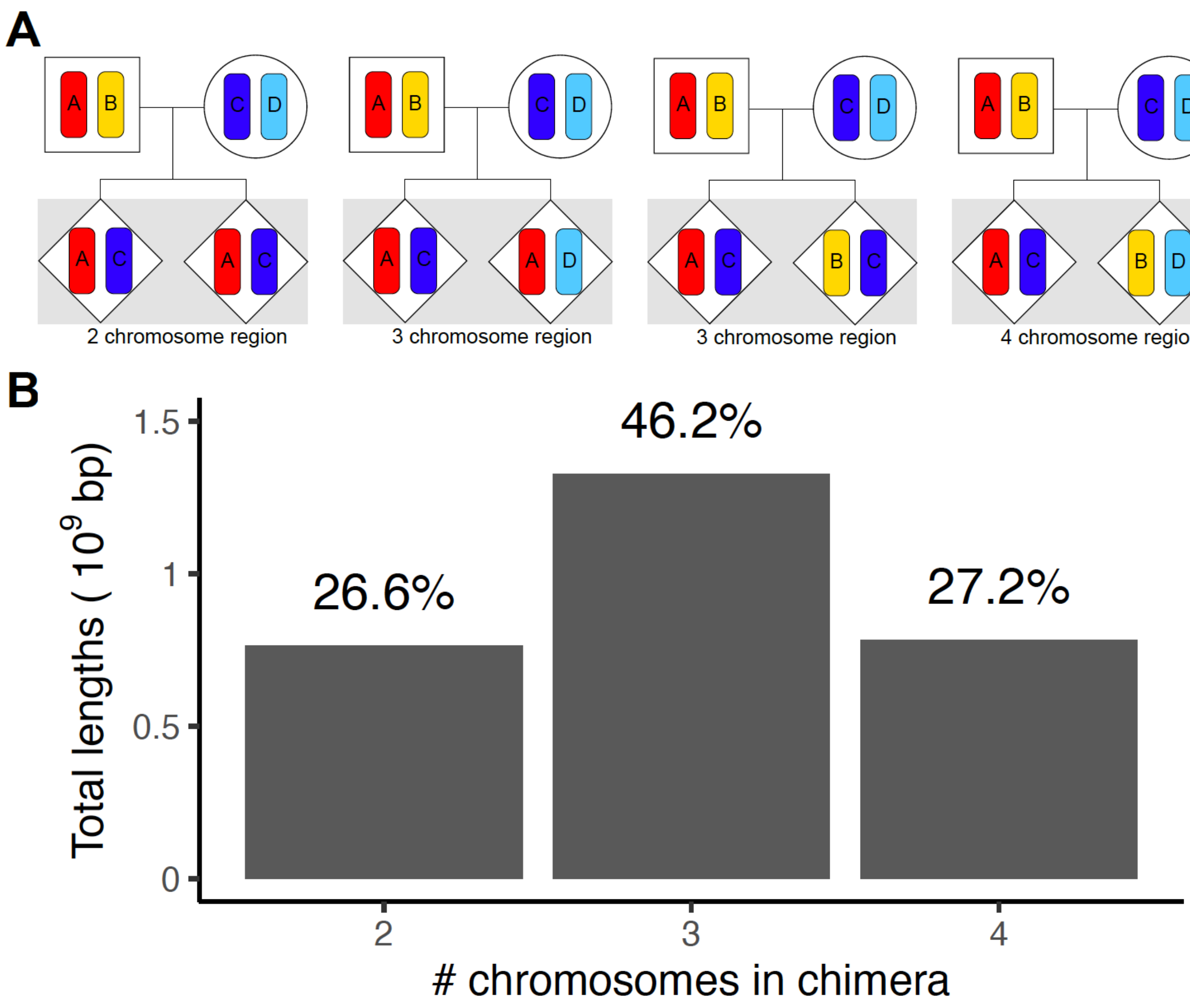


**Supplemental Figure S3. Chromosome distribution of the two zygotes composing the chimera twins.** (A) A possible number of distinct parental chromosomes in the chimera (two zygotes combined, shaded in a gray box) based on the independent inheritance of the chromosomes from the parents. (B) The fraction of genomic regions with each chromatid number within the autosomal genome is shown above each bar plot. The expected distribution of two-chromosome, three-chromosome, and four-chromosome regions from stochastic meiotic recombination in each gamete is 25%, 50%, and 25%.

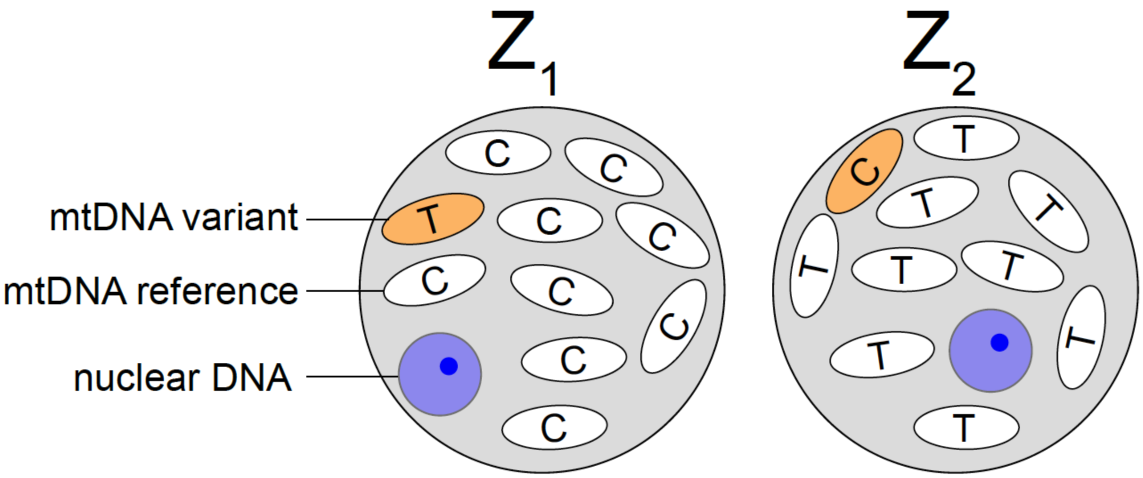


| | | chrM:456 C>T | chrM:13,899 T>C |
|---|---|---|---|
| Child-1 | buccal #1 (60 cells) | 23% | 0% |
| | buccal #2 (60 cells) | 27% | 0% |
| | buccal #3 (60 cells) | 18% | 0% |
| | buccal #4 (60 cells) | 19% | 0% |
| | buccal #5 (60 cells) | 21% | 0% |
| | buccal #6 (25 cells) | 22% | 0% |
| Child-2 | buccal #1 (18 cells) | 0% | 7% |
| | buccal #2 (18 cells) | 0% | 8% |
| | buccal #3 (13 cells) | 0% | 6% |
| | buccal #4 (18 cells) | 0% | 4% |

**Supplemental Figure S4. Single-cell isolated buccal epithelial cell mitochondrial DNA variant analysis.** Mitochondrial heteroplasmy variants unique to each zygote allow tracing of the origins of buccal epithelial cells. LCM-isolated buccal epithelial cells were pooled into 13-60 cells to create a DNA sequence library. Heteroplasmic mitochondrial variant allele fractions for each pooled genetic library are shown.
