## Supplementary material for "Whole-genome sequences reveal zygotic composition in chimeric twins": Key resources table

| REAGENT or RESOURCE | SOURCE | IDENTIFIER |
|---|---|---|
| Antibodies | | |
| Bacterial and virus strains | | |
| Biological samples | | |
| Chemicals, peptides, and recombinant proteins | | |

| | | |
|---|---|---|
| Critical commercial assays | | |
| Carl Zeiss Membrane Slides | Zeiss | Carl Zeiss 415190-9081-000 |
| Zeiss PALM MicroBeam Laser Microdissection | Zeiss | N/A |
| QIAamp DNA micro kit | Qiagen | Cat# 56304 |
| PAXgene Tissue FIX container | PreAnalytiX | Cat#765312 |
| NEBNext ultra II FS DNA library kit | | Cat# E7805 |
| 10x chromium single cell 3' NEXT GEM v3.1 | 10x genomics | PN-1000121 |
| Deposited data | | |
| PBMC single cell RNA seq | This paper | EGAS00001007649 |
| LCM-isolated low input WGS buccal epithelial cell WGS | This paper | EGAS00001007649 |
| Whole genome sequence of buccal and peripheral blood | Yoon et al.[1] | EGAS00001005997 |
| Experimental models: Cell lines | | |
| Experimental models: Organisms/strains | | |

| | | |
|---|---|---|
| Oligonucleotides | | |
| Recombinant DNA | | |
| Software and algorithms | | |
| TrioMix v0.0.2a | Yoon et al.[1] | https://github.com/cjyoon/triomix |
| BWA v 0.7.17-r1188 | Li et al.[2] | https://github.com/lh3/bwa |
| Demuxlet | Kang et al.[3] | https://github.com/statgen/demuxlet |
| CellRanger v6.0.1 | Zheng et al.[4] | https://www.10xgenomics.com/support/software/cell-ranger |
| DNAcopy (R Package) | Seshan et al.[5] | https://bioconductor.org/packages/release/bioc/html/DNAcopy.html |
| Seurat v4.0 | Hao et al.[6] | https://satijalab.org/seurat/ |
| Other | | |

| | | |
|---|---|---|
| Scripts used in this paper | This paper | https://doi.org/10.5281/zenodo.10151688 |
